# Value-related neuronal responses in the human amygdala during observational learning

**DOI:** 10.1101/865568

**Authors:** Tomas G. Aquino, Juri Minxha, Simon Dunne, Ian B. Ross, Adam N. Mamelak, Ueli Rutishauser, John P. O’Doherty

## Abstract

**Abstract:** The amygdala plays an important role in many aspects of social-cognition and reward-learning. Here we aimed to determine whether human amygdala neurons are involved in the computations necessary to implement learning through observation. We performed single-neuron recordings from the amygdalae of human neurosurgical patients (male and female) while they learned about the value of stimuli through observing the outcomes experienced by another agent interacting with those stimuli. We used a detailed computational modeling approach to describe patients’ behavior in the task. Then, using both population-level decoding and single neuron analyses we found evidence to implicate amygdala neurons in two key computations relevant for observational-learning: tracking the expected future reward associated with a given stimulus, and in tracking outcome values received by oneself or other agents. Encoding and decoding analyses suggested observational value coding in amygdala neurons occurred in a different subset of neurons than experiential value coding. Collectively, these findings support a key role for the human amygdala in the computations underlying the capacity for learning through observation.

**Significance statement:** Single neuron studies of the human brain provide a unique window into the computational mechanisms of cognition. In this study, epilepsy patients implanted intracranially with depth microelectrodes performed an observational learning task. We measured activity bilaterally in the amygdala and found a representation for observational rewards as well as observational expected reward values. Additionally, the representation of self-experienced and observational values was performed by distinct subsets of amygdala neurons. This study thus provides a rare glimpse into the role of human amygdala neurons in social cognition.

## Introduction

Acquiring new information about future rewards associated with different stimuli is at the core of an animal’s ability to adapt behavior in order to maximize net future rewards (Sutton and Barto, 2018). In many organisms, reinforcement learning (RL) can take place not only through taking actions and experiencing outcomes, but also indirectly through observing the actions taken and outcomes obtained by others, in a form of learning known as observational learning (OL) (Cooper et al., 2012; Van Den Bos et al., 2013; Dunne et al., 2016; Charpentier and O’Doherty, 2018). The computational and neural basis of reinforcement-learning through direct experience has been the focus of intense study, and as a result much is now known about its neural underpinnings (Doya, 1999; Daw et al., 2005; Lee et al., 2012; O’Doherty et al., 2017). In contrast, the neural mechanisms of learning through observation has been much less well studied, especially in humans.

A core feature of RL models is that in order to decide whether or not to choose a particular stimulus or action, it is first necessary to consider the expected future reward associated with that option. Consistent with this notion, neuronal activity has been found in the amygdala as well as elsewhere in the brain, which tracks the expected future reward associated with various options at the time of decision-making (Gottfried et al., 2003; Holland and Gallagher, 2004; Hampton et al., 2006; Salzman and Fusi, 2010; Wang et al., 2017). Lesions of the amygdala have shown it is necessary for guiding behavior on the basis of expected future outcomes learned about through experience (Málková et al., 1997; Bechara et al., 1999; Schoenbaum et al., 2003; Hampton et al., 2007; De Martino et al., 2010), suggesting that value representations in this area are causally relevant for driving value-related behavior. The amygdala performs these functions in concert with a broader network that regulates reward learning, memory and emotion (Murray, 2007). For instance, adaptive responses to reward cues and devaluation depend on amygdala-OFC connections in monkeys (Baxter et al., 2000) and mice (Lichtenberg et al., 2017). The amygdala also receives significant dopaminergic projections from the ventral tegmental area (VTA) and the substantia nigra pars compacta (SNc) (Aggleton et al., 1980).

Turning to the role of the amygdala in observational learning specifically, recent evidence has suggested a role for amygdala neurons in non-human primates in responding to the rewards obtained by others (Chang et al., 2015) as well as to others’ choices (Grabenhorst et al., 2019). However, much less is known about the role of the amygdala in the human brain in processes related to OL. One human single neuron study reported amygdala neurons which tracked observational outcomes (Hill et al., 2016), though the most robust signals were found in rACC. Building on evidence implicating the human amygdala not only in reward processing but also in social cognition more broadly (Rutishauser et al., 2015a), we aimed to address the extent to which neurons in the human amygdala are specifically involved in contributing to observational learning. For this, we administered an observational learning task to a group of human neurosurgery patients while obtaining single-neuron recordings from electrodes located in the amygdala. This provided us with a rare opportunity to investigate the role of amygdala neurons in value prediction coding and updating during OL. We hypothesized that we would find evidence for reinforcement-learning signals in the amygdala during observational learning, especially concerning the representation of the value of stimuli learned through observation. Furthermore, we also aimed to compare and contrast the contribution of amygdala neurons to OL with that of the role of these neurons in experiential learning. Of particular interest was the question of whether an overlapping or distinct population of neurons in the amygdala contributes to encoding reinforcement-learning variables in observational compared to experiential learning.

## Materials and Methods

### Electrophysiology and electrodes

Broadband extracellular recordings were filtered from 0.1Hz to 9kHz and sampled at 32 kHz (Neuralynx Inc). The data reported here was recorded bilaterally from the amygdala, with one macroelectrode in each side. Each of these macroelectrodes contained eight 40 *μm* microelectrodes. Recordings were performed bipolar, with one microwire in each area serving as a local reference (Minxha et al., 2018). Electrode locations were chosen exclusively according to clinical criteria.

### Patients

Twelve patients (4 females) who were implanted with depth electrodes prior to possible surgical treatment of drug resistant localization related epilepsy volunteered to participate and gave informed consent. Four of the patients performed two recording sessions, and the others performed only one. One pilot session was not included in the analysis and one session was discarded due to technical error. Protocols were approved by the Institutional Review Boards of the California Institute of Technology, the Cedars-Sinai Medical Center and the Huntington Memorial Hospital. Electrode localization was determined based on pre and post-operative T1 scans, using previously published methods (Kamiński et al., 2017) and the CIT168 template brain (Tyszka and Pauli, 2016), registered on MNI152 space (Fig. 2d).

### Spike detection and sorting

Spike detection and sorting was performed as previously described using the semiautomatic template-matching algorithm OSort (Rutishauser et al., 2006). Channels with interictal epileptic activity were excluded. Across all valid sessions, we isolated in total 202 putative single units in amygdala. We will refer to these putative single units as “neuron” and “cell” interchangeably. Units isolated from electrodes localized outside of the intended target region were not included in the analyses. We characterized the quality of the isolated units using the following metrics: the percentage of interspike intervals (ISIs) below 3 ms was 0.49% ± 0.63%; the mean firing rate was 1.98*Hz* ± 2.47*Hz*; the SNR at the mean waveform peak, across neurons, was 5.12 ± 3.24; the SNR of the mean waveform across neurons was 1.87 ± 0.97; the modified coefficient of variation (CV2) (Holt et al., 1996) was 0.95 ± 0.11; and the isolation distance (Schmitzer-Torbert et al., 2005) was 1.69 ± 0.59 for neurons in which it was defined.

### Task and behavior

Patients performed a multi-armed bandit task (Dunne et al., 2016) with 288 trials in total, distributed across 2 experiential and 2 observational blocks. Block order was chosen to always interleave block types, and the type of the initial block was chosen randomly (see Fig. 1a). Each block had 72 trials, out of which 48 were no-choice trials and 24 were binary choice trials. Experiential no-choice trials began with the presentation of a single bandit, whose lever was pulled automatically 0.5s after stimulus onset. Each block consisted of two possible bandits that were chosen randomly in every trial. Subjects were told that the color of a bandit allows them to differentiate the different bandits. Bandits were repeated across blocks of the same type, with the possibility of contingency reversal. Outcome was presented 1s after the automatic lever press, and subjects received feedback on the amount of points won or lost in the trial, which was added or subtracted to their personal total. The amount of points for each trial was selected from a normal distribution, with specific means and variances for each bandit, truncating at −50 and +50 points. Observational no-choice trials consisted in watching a pre-recorded video of another player experiencing the same trial structure. Points received by the other player in the pre-recorded video were not added or subtracted to subjects’ personal total, and subjects were informed of this fact. Choice trials started with the presentation of two bandits, and subjects had up to 20s to select one via button press, which would cause the lever on the corresponding bandit to be pulled. If subjects failed to respond within 20s, the trial was considered missed, and subjects received a penalty of 20 points. In choice trials, after a 1s period, subjects observed closed curtains on the screen instead of outcome feedback. Subjects were told they would still receive or lose the amount of points displayed behind the curtain, despite the lack of feedback. This was done to restrict learning to no-choice trials, to further dissociate the decision making and reward learning components of the task. The two bandits that could be chosen in choice trials were always the two possible bandits from no-choice trials in the current block. Intertrial intervals were jittered with a uniform distribution between 1s and 3s regardless of block and trial type. To further motivate subjects, a leaderboard was shown in the end of the task, displaying the amount of points won by the subject, in comparison to amounts won by previous participants.

**Figure 1:**
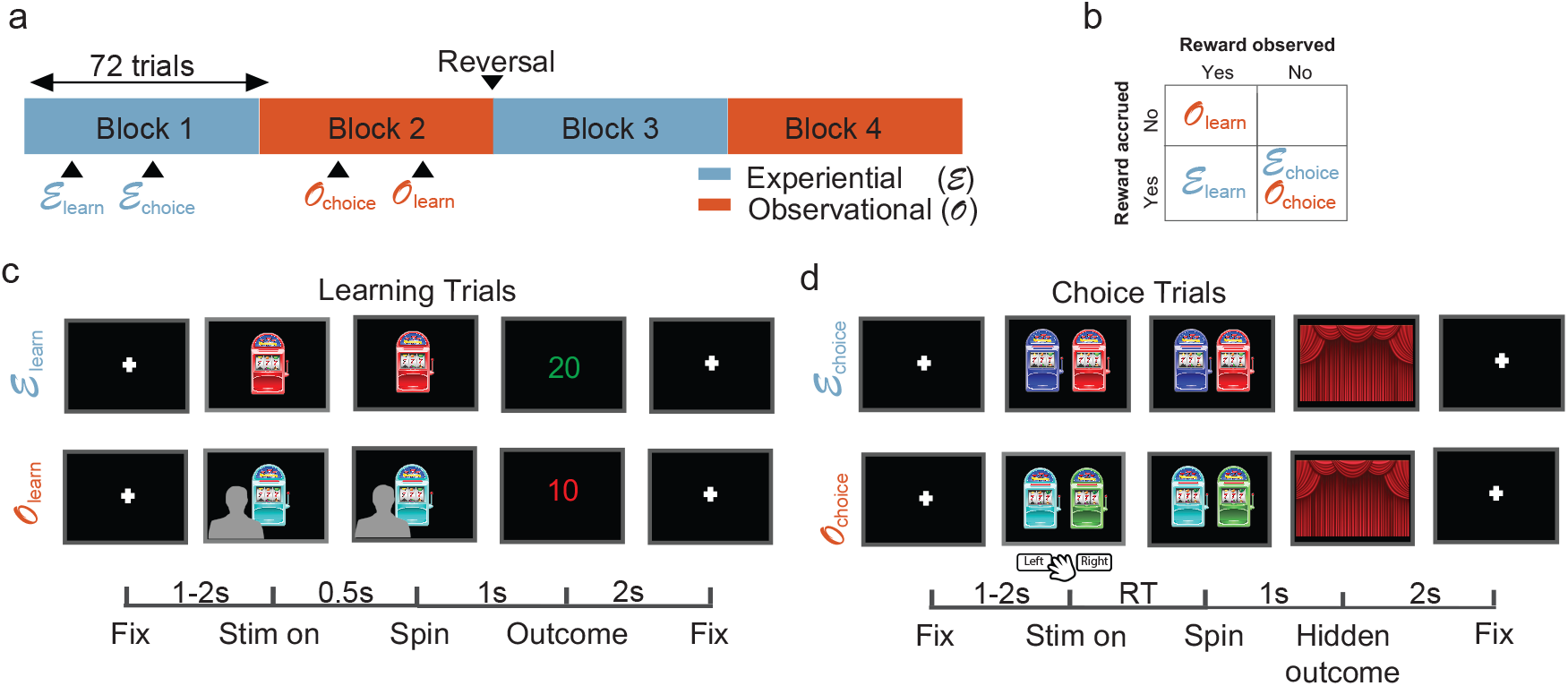
Observational learning task. (a) Block structure. The task had 288 trials in total, in 4 blocks of 72 trials. Each block contained either experiential or observational learning trials, as well as choice trials. Block order was interleaved, and bandit values were reversed after the end of block 2. (b) Reward structure. Reward was accrued to subjects’ total only in experiential trials, and reward feedback was only presented in learning trials, both in experiential and observational blocks. (c) Learning trials structure. Top row: experiential learning trials. After a fixation cross of jittered duration between 1-2s, subjects viewed an one-armed bandit whose tumbler was spun after 0.5s. After a 1s spinning animation, subjects received outcome feedback, which lasted for 2s. Bottom row: observational learning trials. Subjects observed a video of another player experiencing learning trials with the same structure. Critically, outcomes received by the other player were not added to the subject’s total. Lower bar: timing of trial events in seconds. (d) Choice trials structure. Subjects chose between the two bandits shown in the learning trials of the current block. After deciding, the chosen bandit’s tumbler spun for 1s, and no outcome feedback was presented.

**Figure 2:**
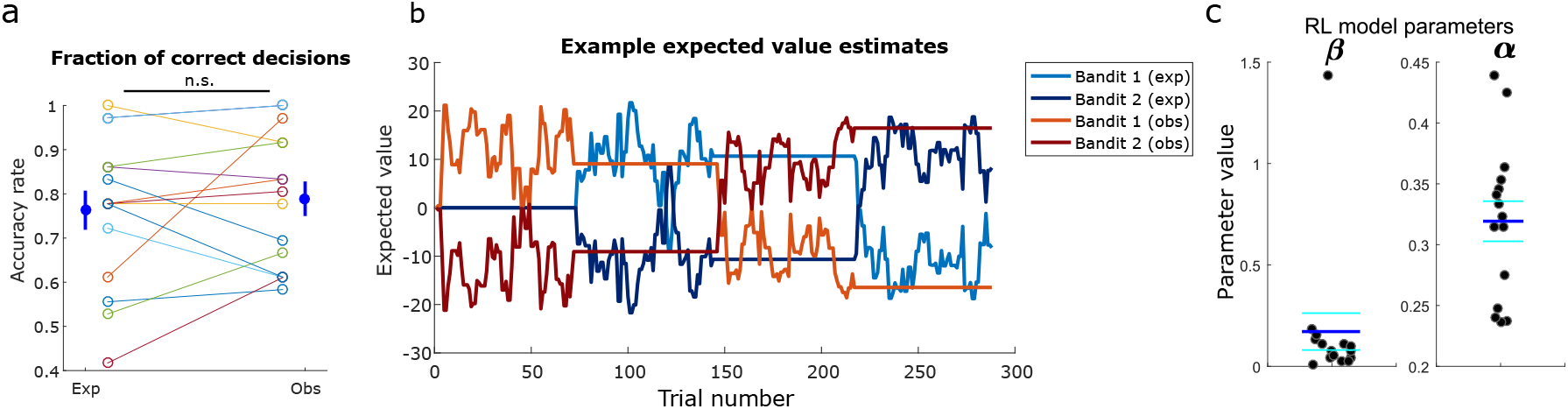
Behavior and reinforcement learning model. (a) Accuracy rate of all sessions as defined by the fraction of free trials in which a subject chose the bandit with highest mean payout, discarding the first 25% of trials in each block. Each color represents a different session, for experiential and observational trials, with average and standard error indicated on the left and right. Accuracy in experiential and observational trials was not significantly different (p < 0.66, two-sample t-test). (b) Typical time course of modeled EVs throughout the task, using the RL (counterfactual) model. Bandit 1 (exp) and Bandit 2 (exp) indicate EVs for each of the two bandits shown in experiential blocks respectively, whereas Bandit 1 (obs) and Bandit 2 (obs) indicate EVs for each of the two bandits shown in observational blocks, respectively. (c) Parameter fits for each valid session, for the chosen reinforcement learning model. The model contained a single learning rate (*α*)for experiential and observational trials, and an inverse temperature *β*. Dark blue horizontal lines indicate parameter means and cyan horizontal lines indicate standard error.

### Computational modeling

We focused on the form of observational learning referred to as vicarious learning, which takes place when individuals observe others taking actions and experiencing outcomes, rather than doing so themselves (Charpentier and O’Doherty, 2018). At the computational level, we hypothesized that vicarious learning involves similar mechanisms to those utilized for experiential learning. To test this hypothesis, we adapted a simple model-free learning algorithm from the experiential to the observational domain (Cooper et al., 2012). For both observational and experiential learning, this model learns EVs for each stimulus via a reward prediction error (RPE) that quantifies discrepancies between predicted values and experienced reward outcomes. This prediction error signal is then used to iteratively update the value predictions.

We used the behavior in the choice trials to fit four different types of computational models. We used a hierarchical Bayesian inference framework to achieve both hierarchical model fitting and model comparison (Piray et al., 2019). This framework allowed us to infer a protected exceedance probability for each model, as well as individualized model parameters for each subject. The model with the largest protected exceedance probability was chosen for the model-based encoding and decoding analyses.

We first provide a brief summary of each of the computational models before describing each in detail. The first model was a simple reinforcement learning model (Sutton and Barto, 2018) with a single learning rate parameter for both experiential and observational trials (RL (no split)); the second model was the same, except that learning rates were split between observational and experiential trials (RL (split)); the third model was a counterfactual reinforcement learning model with a single learning rate in which EVs for played bandits were updated as usual, but EVs for the bandits that were not seen in a trial were also updated, in the opposite direction of the bandits that were actually played (RL (counterfactual)). The last model was a hidden Markov model with built-in reversals, with two states. The first state assumed one of the bandits in the block had a positive mean payout, while the other bandit had a negative mean payout with the same magnitude. The second state mirrored the first one, switching which bandits had the positive and negative payouts. This model allowed us to include inferred reversals between those two states, and to model the inferred reversal rate that patients assumed to be true. Expected values in all models were initialized to zero for all bandits.

The RL (no split) model keeps a cached value V for the EV of each bandit i, in every trial t, updated according to the following rule:

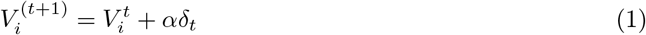

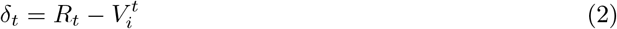

In this case, *α* represents the learning rate for both the experiential and observational cases, *δ* represents reward prediction error (RPE) and R represents reward feedback value. The RL (split) model is identical, except that a learning rate *α_exp_* is applied in experiential trials and another learning rate *α*_*obs*_ is applied in observational trials.

The RL (counterfactual) model is identical to RL (no split), except that both bandits are updated on every trial, in opposite directions. For the chosen and unchosen bandits in every trial, the EV updates are as follows:

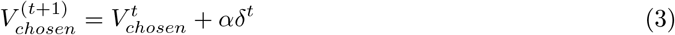

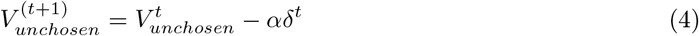

The HMM has been formalized similarly to previous work (Prévost et al., 2013). An inferred state variable *S*_*t*_ represented the association between bandits and rewards at trial *t*. Assuming the two bandits in a block are arbitrarily indexed as A and B, that the magnitude of the inferred mean payout was a free parameter *μ* fixed throughout the task, and that *mu*_*A*_ and *mu*_*B*_ denote the mean payouts for bandits A and B respectively,

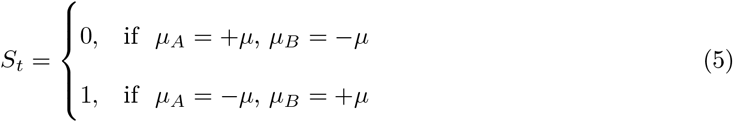

This model allows for inferring reversals between states, which means the inferred mean payouts of the two bandits are swapped. The reversal structure is dictated by the following reversal matrix, assuming reversal rates were a free parameter *r* fixed throughout the task:

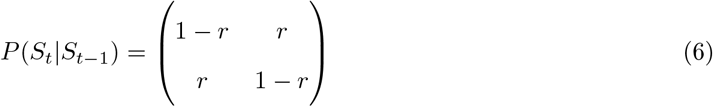

Given this transition structure, the prior *Prior*(*S*_*t*_) would be updated in every trial as follows:

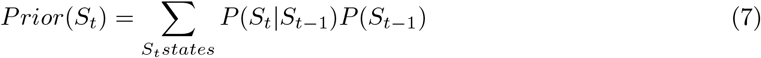

Initial state probabilities were set to 0.5. Then, using Bayes’ rule, the posterior would be updated using evidence from the outcome *R*_*t*_:

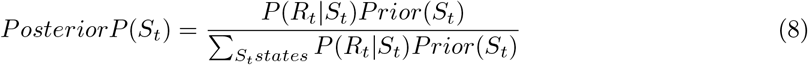

Outcome variables were assumed to have a Gaussian distribution, with a fixed standard deviation free parameter *σ*:

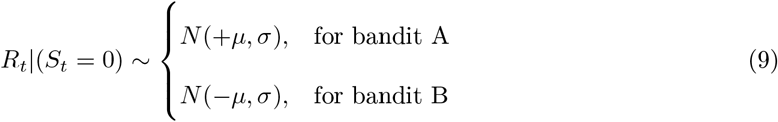

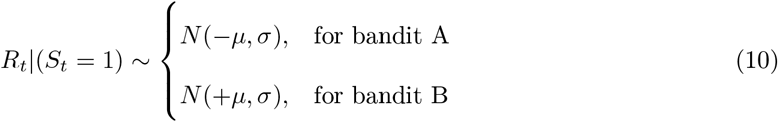

This framework allowed for computing EVs in each trial *t* for each bandit *j*, taking into account the probability of being in each state:

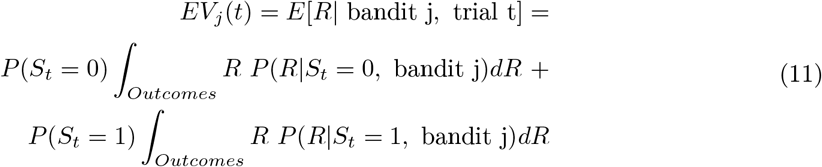

Since outcomes were assumed to be normally distributed, for each bandit this reduced to

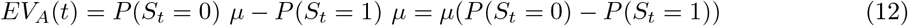

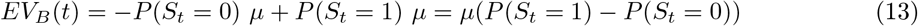

This means that EVs for a certain bandit were larger if patients inferred they were more likely in the state in which that bandit was better. For example, if *P*(*S*_*t*_ = 0) = 0.9 and *P*(*S*_*t*_ = 1) = 0.1, then:

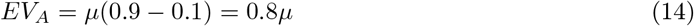

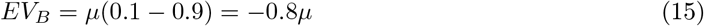

We used the cached value estimates of EV as parameters in a softmax function controlled by an inverse-temperature parameter *β* for each session, to generate decision probabilities in free-choice trials. For the RL models, we constrained *α*, *α*_*exp*_, and *α*_*obs*_ in the (0,1) interval, and *β* in the (0,10) interval. In the HMM, we constrained r in the (0,1) interval, and both *μ* and *σ* in the (0,20) interval.

Model comparison was performed by computing the protected exceedance probability of each model and selecting the one with the largest value, which was RL (counterfactual) (Fig. 3). For all subsequent model-based analyses, we display results using the EVs and RPEs produced by the RL (counterfactual) model. An example of how the EVs assigned to each bandit typically behave in a modeled session is displayed in Fig. 2b.

**Figure 3:**
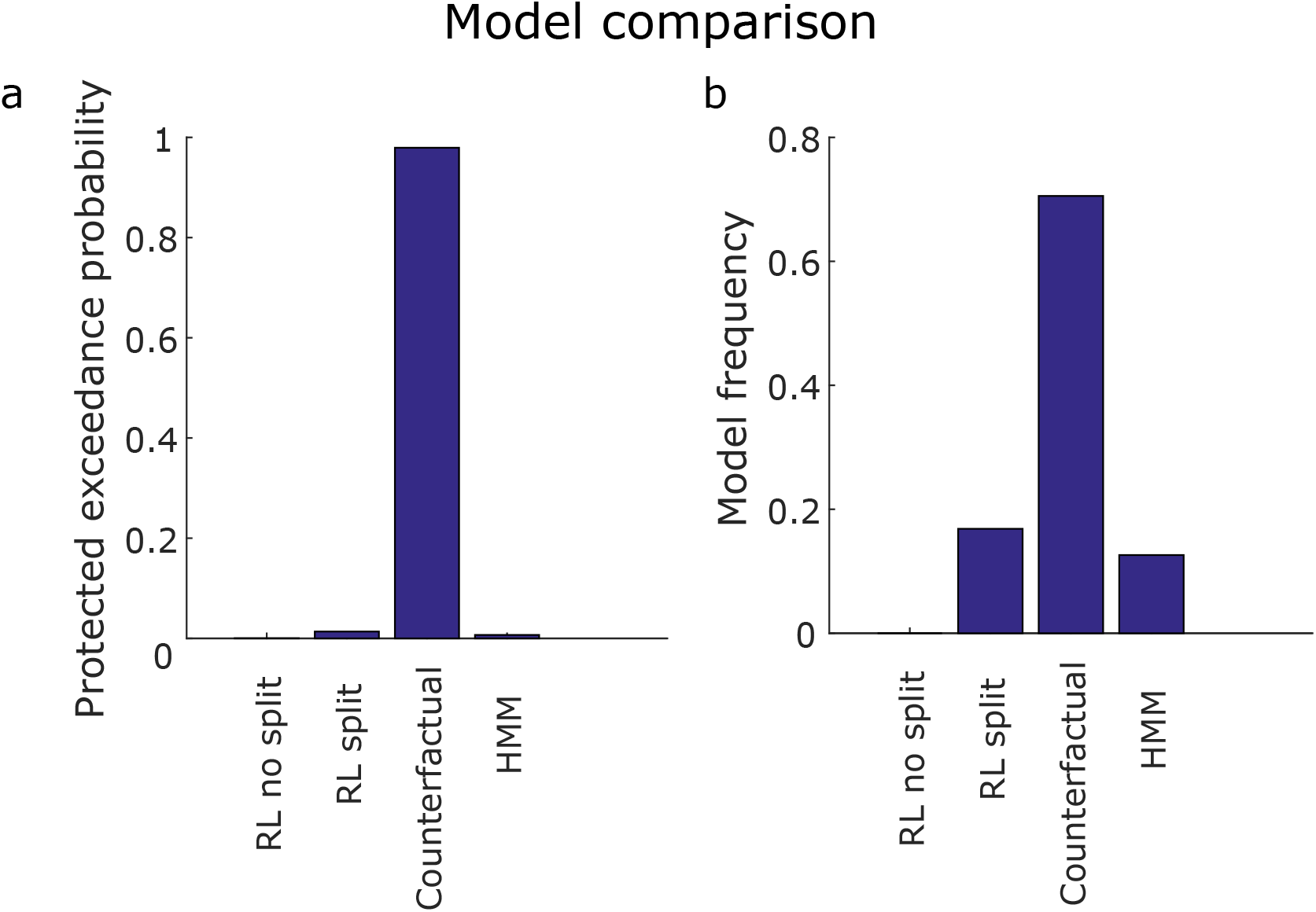
Model comparison. (a) Protected exceedance probability. This is the probability that each one of the four models fit using hierarchical Bayesian inference (RL split, RL no split, Counterfactual, and HMM) was more likely than any other, taking into account the possibility that there is no difference between models. (b) Model frequency. This is the proportion of individual patients whose behavior is better explained by each model. The counterfactual learning model outperforms the others both in terms of protected exceedance probability and model frequency.

### Population decoding analysis

Population decoding was performed with the Neural Decoding Toolbox (Meyers, 2013) as described previously (Rutishauser et al., 2015b). We pooled neurons from all sessions into a single pseudopopulation with 202 amygdala neurons. To achieve this alignment on a trial by trial basis across sessions, we created discrete trial bins using quantiles of the decoded variable, with the same number of trials, for each session. For example, for outcome decoding, we found which trials for each session fit into each one of 4 quantiles of received outcomes and aligned trials that fell in the same bin across sessions, assuming all neurons belonged to the same session. This session-based trial binning meant the exact quantile boundaries were not necessarily the same across sessions. For example, a trial in which the outcome was 10 points might have been placed on bin 2 for one session and on bin 3 for another session, depending on the distribution of outcomes in each session. Finally, all learning trials across sessions had the same event timing, so no additional temporal alignment was needed.

We used this strategy to create a neural activity tensor of dimensions (*n*_*neurons*_, *n*_*trials*_), where *n*_*trials*_ is the number of trials in a single session. Decoding consisted of training and testing a classifier tasked with correctly predicting which bin of the variable of interest each trial belonged to, only from information contained in the neural activity tensor.

We used a maximum Pearson correlation classifier with access to spike counts binned in a time window of interest. This classifier learns by obtaining a mean representation *x*_*c*_ of each class *c* in the multidimensional neural population space, and assigns new data points *y* to a class *c*\* corresponding to *c*\* = *argmax*_*c*_(*corr*(*x*_*c*_, *y*)). We used 10-fold cross-validation and 20 cross-validation fold resample runs, which were averaged to generate a testing decoding accuracy score. Significance was determined via permutation test with 500 re-runs, shuffling regressor labels. Expected value decoding was only tested in the pre-outcome period (300ms to 1500ms from trial onset), whereas outcome and prediction error decoding was only tested in the post-outcome period (300 to 2000ms from outcome onset).

### Single neuron encoding analysis

For every tested neuron *n*, we used a Kruskal-Wallis test (Kruskal and Wallis, 1952) to fit binned spike counts *y*_*n*_(*t*) (1200 ms bins for pre-outcome, 1700 ms bins for post-outcome, 3500ms bins for the whole trial), implemented with the MATLAB function *kruskalwallis*. Outcome and prediction errors were regressed only on the post-outcome period, and expected values were regressed only on the pre-outcome period. Trial type regression was performed in the entire trial. Significance was determined through permutation tests by shuffling variable labels. For expected value and prediction error time series, in which trials might not be independent from each other, we performed variable shuffling using surrogate time series as described previously (Schreiber and Schmitz, 2000). For the other variables, we used standard random permutations. We then used chi-squares yielded by the Kruskal-Wallis test as a statistic for each regressor.

## Results

### Behavioral performance

We obtained a behavioral metric of subject performance on choice trials (Fig. 2a): we defined “correct” trials as those in which the subject selected the bandit with the highest mean payout, disregarding the first 25% of trials in each block, to exclude the confounding effects of initial learning. We found that in 12 out of 15 recording sessions, performance was above the 95% percentile of a random agent, theoretically determined by a binomial distribution with success probability 0.5, thereby indicating that behavior in these sessions was significantly better than chance.

Overall, subjects performed well in both the experiential and observational condition (Fig. 2a): The average correct choice ratio in all trials was 0.776 ± 0.038. Taking only experiential trials, the average correct choice ratio was 0.763±0.044; in observational trials, it was 0.789±0.039. Experiential and observational correct choice ratios were not significantly different from each other across all sessions (two sample t-test, p = 0.6641 > 0.05).

### Computational model fitting

We fit four computational models to each subjects’ behavior during choice trials (see methods): a model-free reinforcement learning model with one learning rate for experiential and observational trials (RL (no split)); a model-free reinforcement learning model with separate learning rates split between experiential and observational trials (RL (split)); a counterfactual reinforcement learning model in which outcomes from the played bandit also were used to update EVs for the unseen bandit in each trial; and a hidden Markov model (HMM) with an estimate of reversal rates on a trial-by-trial basis. In all models, we applied a softmax rule to generate probabilistic decisions. Model fitting and comparison were performed simultaneously with hierarchical Bayesian inference (HBI) (Piray et al., 2019), described in more detail in the methods section.

Overall, the counterfactual RL model, with a single learning rate for experiential and observational trials, outperformed the others in both protected exceedance probability (Fig. 3a) and inferred model frequency among the patient population (Fig. 3b). The mean learning rate in the winning model was 0.31 ± 0.06, and the mean softmax inverse temperature *β* was 0.17 ± 0.35 (Fig. 2c). Taken together, these behavioral findings suggest subjects employed a similar learning strategy for the valuation of each bandit regardless of trial type, and were still engaged with the task when another person received rewards. Given that the counterfactual model was the best fitting model for explaining participants’ behavior on the task, we utilized the variables generated by this model in the subsequent computational model-based analysis of the neuronal data.

### Amygdala population decoding

We tested whether the activity of amygdala neurons was related to the following task and computational variables: trial type, EV, outcome, and RPE, during learning trials. Trial type decoding was performed in the whole trial; EV decoding was performed in a 1200ms time bin starting 300 ms from stimulus onset, until outcome presentation; outcome and RPE decoding was performed in a 1700ms time bin starting 300ms after outcome presentation.

For each variable, we trained a maximum Pearson correlation classifier on a pseudopopulation of amygdala neurons (see methods; Fig. 4). Cross-validated decoding accuracy was obtained for each tested variable, tested for significance through a permutation test with 500 shuffled label runs. The same procedure was repeated in 50 cross-validation randomly re-sampled folds. To perform decoding of continuous variables across sessions, we binned variables (EV, outcome, and RPE) into 4 bins (quantiles). P-values were obtained by computing the proportion of shuffled instances in which decoding accuracy exceeded the real decoding accuracy. With this method, the smallest p-value attainable was 1/*n*_*permutations*_ = 0.002.

**Figure 4:**
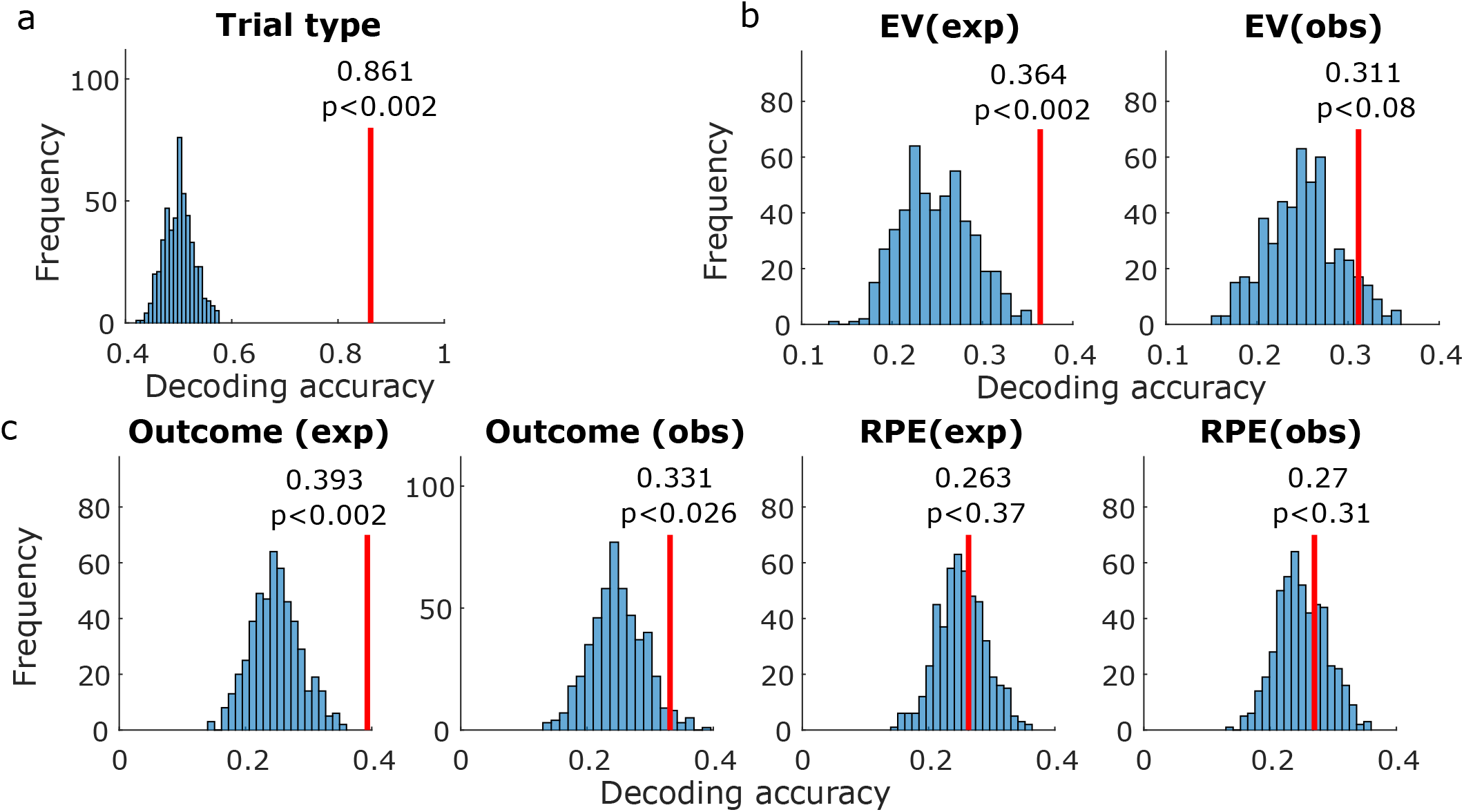
Amygdala population decoding analysis. (a) Entire trial decoding. The tested variable was trial type (experiential vs. observational). The vertical red line indicates average decoding accuracy in held-out trials after training with a maximum Pearson correlation classifier. The histogram indicates decoding accuracy in each instance of a permutation test, shuffling variable labels. P-values were obtained by computing the proportion of permutation iterations in which the decoding accuracy exceeded the true decoding accuracy. (b) Same, decoding within the pre-outcome period. Decoded variables, from left to right, were EV (experiential) and EV (observational). (c) Same, decoding within the post-outcome period. Decoded variables, from left to right, were outcome (experiential), outcome (observational), RPE (experiential), and RPE (observational).

#### Trial type decoding

Trial type (experiential vs observational) could be decoded from amygdala neurons with above chance accuracy (see Fig. 4a; *p* < 0.002 < 0.05, permutation test). Average decoding accuracy in held-out trials was 86.1%. This indicates that amygdala neurons prominently tracked whether the current block was experiential or observational.

#### Expected value decoding

We next tested whether EV was decodable in the pre-outcome period (300ms to 1500ms from bandit onset), separately for observational and experiential learning trials. We found better than chance decoding in experiential trials (see Fig. 4b; *p* < 0.002 < 0.05, permutation test). Average experiential EV decoding accuracy in the pre-outcome period was 36.4%. In contrast, observational EV decoding was within the chance boundaries of the permutation test (*p* < 0.08). (Fig. 4b). This indicates that amygdala neuron populations contained more easily decodable information for keeping track of rewards received by oneself than by the other player.

#### Outcome decoding

Following outcome onset (300ms to 2000ms from outcome onset), outcome was decodable above chance in experiential trials (see Fig. 4c; *p* < 0.002 < 0.05, permutation test), with an average decoding accuracy of 39.3%. Additionally, outcome was also decodable above chance in observational trials (see Fig. 4c; *p* < 0.026 < 0.05, permutation test), with an average decoding accuracy of 33.1%. This indicates that amygdala populations represented both experienced and observed outcomes, but more strongly in the experienced case.

#### Reward prediction error (RPE) decoding

We tested for decodability of RPEs during the outcome period (300ms to 2000ms from outcome onset), but did not find better decoding accuracy than expected by chance in the permutation test (see Fig. 4c), both in the experiential (*p* < 0.37, permutation test) and the observational cases (*p* < 0.31, permutation test).

### Single neuron encoding analysis

In order to understand the relationship between the population decoding result and the activity of single neurons we next tested the sensitivity of each amygdala neuron (*n* = 202 neurons) to each one of the decoded variables (Fig. 5). We used a Kruskal-Wallis analysis to compare every individual neuron’s activity to the same variables used in decoding. We chose this method as opposed to a GLM analysis to encompass units whose activities might be non-linearly modulated by a variable of interest (e.g. being less active for intermediate levels of a variable of interest), such as the one displayed in (Fig. 6d).

**Figure 5:**
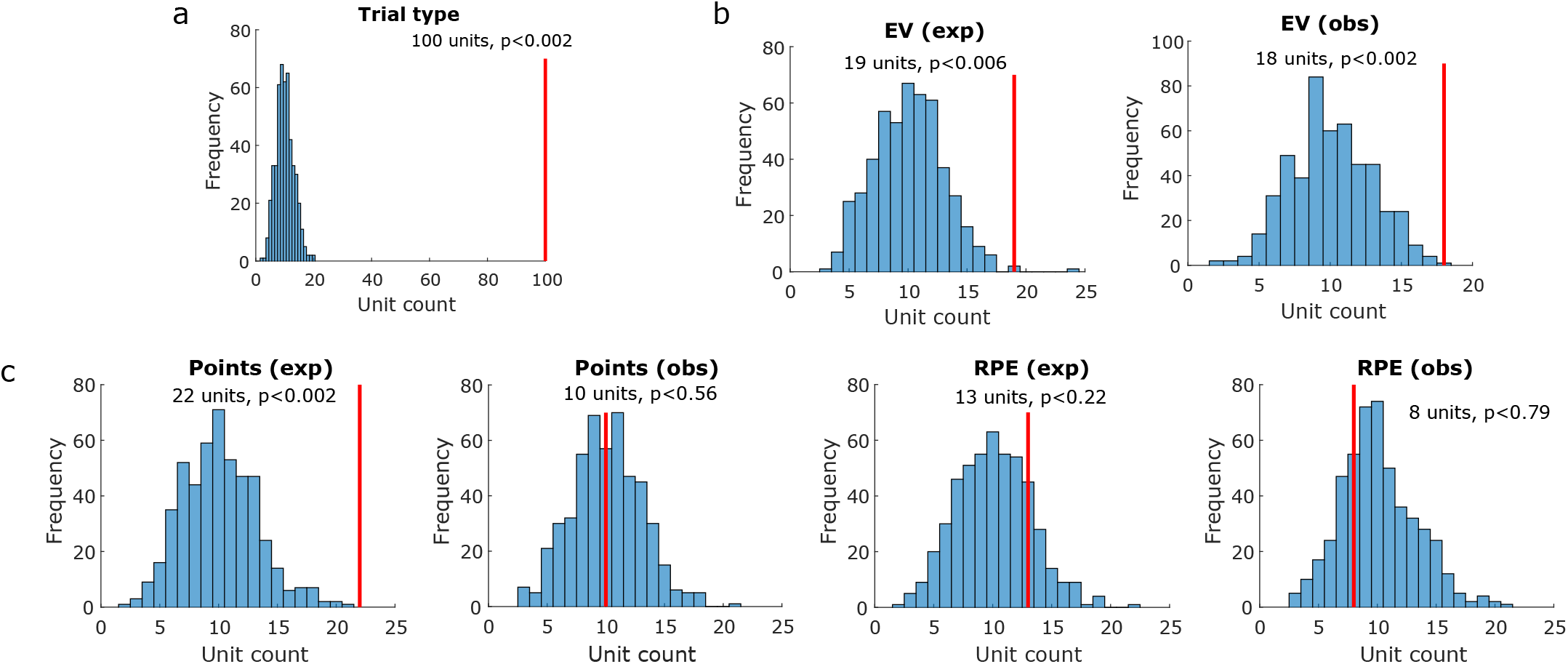
Amygdala single neuron encoding analysis. (a) Whole-trial encoding of trial type (experiential vs. observational). Vertical red lines indicate how many units were found to be sensitive to the tested variable within the pre-outcome period. Histograms indicate how many units were sensitive to the tested variable in each iteration of the permutation test, shuffling variable labels. P-values were obtained by computing the proportion of permutation iterations in which unit counts exceeded the true unit count. (b) Pre-outcome encoding of EV in experiential (left) and observational (right) trials. (c) Same, but for the post-outcome period. Encoded variables, from left to right, were outcome (experiential), outcome (observational), RPE (experiential), and RPE (observational).

**Figure 6:**
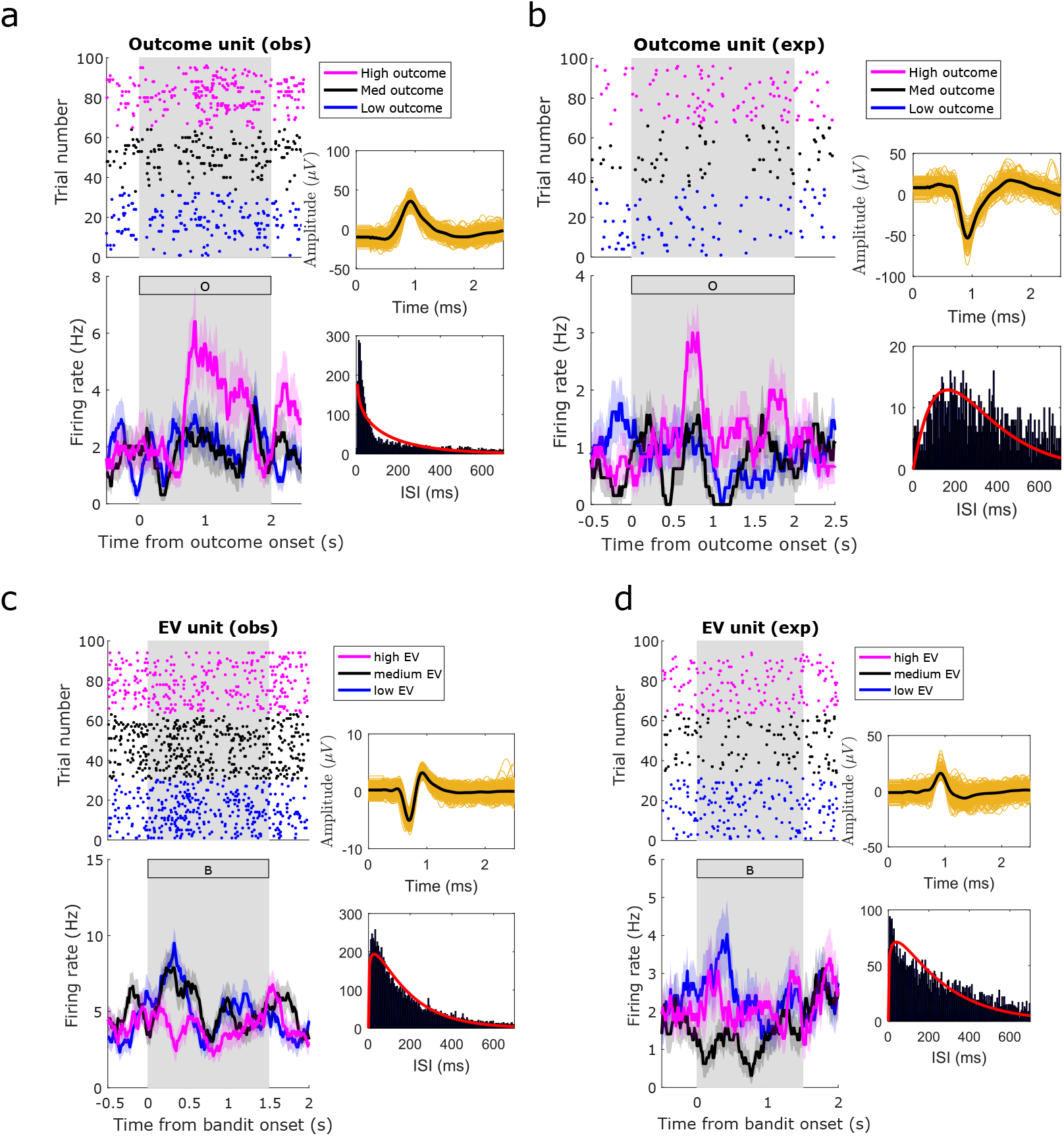
Amygdala neuron raster plot examples. Example amygdala units, significantly modulated by the indicated regressors, in the indicated conditions. (a) Unit modulated by outcome in observational trials during post-outcome period. (b) Unit modulated by outcome in experiential trials during post-outcome period. (c) Unit modulated by EV in observational trials during pre-outcome period. (d) Unit modulated by EV in experiential trials during pre-outcome period. Top: raster plots. For plotting purposes only, we reordered trials by regressor levels by obtaining 3 quantiles from the variable of interest (magenta: high; black: medium; blue: low). Bottom: PSTH (bin size = 0.2*s*, step size = 0.0625*s*). The annex panels to the right of each raster display spike waveforms (top) and interspike interval histograms (bottom) from the plotted neuron. Background gray rectangles post-outcome periods, for (a) and (b), or pre-outcome periods, for (c) and (d). Rectangles filled with a letter indicate which stimulus was present on the screen at that time (B: bandit; O: outcome).

#### Trial type neurons

We found 100 amygdala neurons whose activity is significantly different across experiential and observational trials (49.5%, *p* < 0.002 < 0.05, permutation test). Note this could partially be explained as an effect of the blocked design we chose, grouping all experiential trials and observational trials in distinct trial blocks. This result is also consistent with the high trial type decoding accuracy we found in left-out trials.

#### Expected value neurons

We tested amygdala neurons for experiential EV sensitivity, and found 19 sensitive units (9.4%, *p* < 0.006 < 0.05, permutation test) during the pre-outcome period (Fig. 5b, left). One experiential EV example unit is shown in Fig. 5d. Conversely, observational EV sensitivity was found in 18 units (8.9%, *p* < 0.002 < 0.05, permutation test). An observational EV example unit is shown in Fig. 5c. Taken together, these findings suggest that the expectation of outcomes is represented in a significant proportion of amygdala neurons, both for experienced and observed outcomes.

#### Outcome neurons

We also tested amygdala neurons for outcome sensitivity, in the post-outcome period. For experiential outcomes, we found a significant proportion (10.8%, *p* < 0.002 < 0.05, permutation test) of sensitive amygdala neurons (Fig. 5c, first panel). For observational outcomes (Fig. 5c, second panel), however, only 10 units were selected as sensitive (4.9%, *p* < 0.56, permutation test), despite better-than-chance observational outcome decoding. Example outcome neurons are displayed in Fig. 6a (observational) and Fig. 6b (experiential).

#### Reward prediction error neurons

Also in the post outcome period, we found 13 (6.4%, *p* < 0.22, permutation test) experiential RPE units (Fig. 5c, third panel), as well as 8 (3.9%, *p* < 0.76, permutation test) observational RPE units (Fig. 5c, fourth panel). Neither of these unit counts exceeded what is expected by chance in the permutation test. This finding is consistent with the low decoding accuracy we obtained for reward prediction errors in the population decoding analyses.

### Decoding generalization analysis

To test whether the same or different neurons encode experienced and observational variables we performed a decoding generalization analysis (Wang et al., 2019). We trained decoders with neural activity in experiential trials and tested in observational trials (Fig. 7a), and vice-versa (Fig. 7b). The method is otherwise identical to the previous decoding analysis. We tested generalization of EVs (Fig. 7a,b, left panels) in the pre-outcome period and outcomes in the post-outcome period (Fig. 7a,b, right panels), since these variables were represented in the amygdala neuron population to some extent: outcome decoding was successful in both trial types, and despite weaker observational EV decoding, we did find a significant observational EV unit count through the encoding analysis.

**Figure 7:**
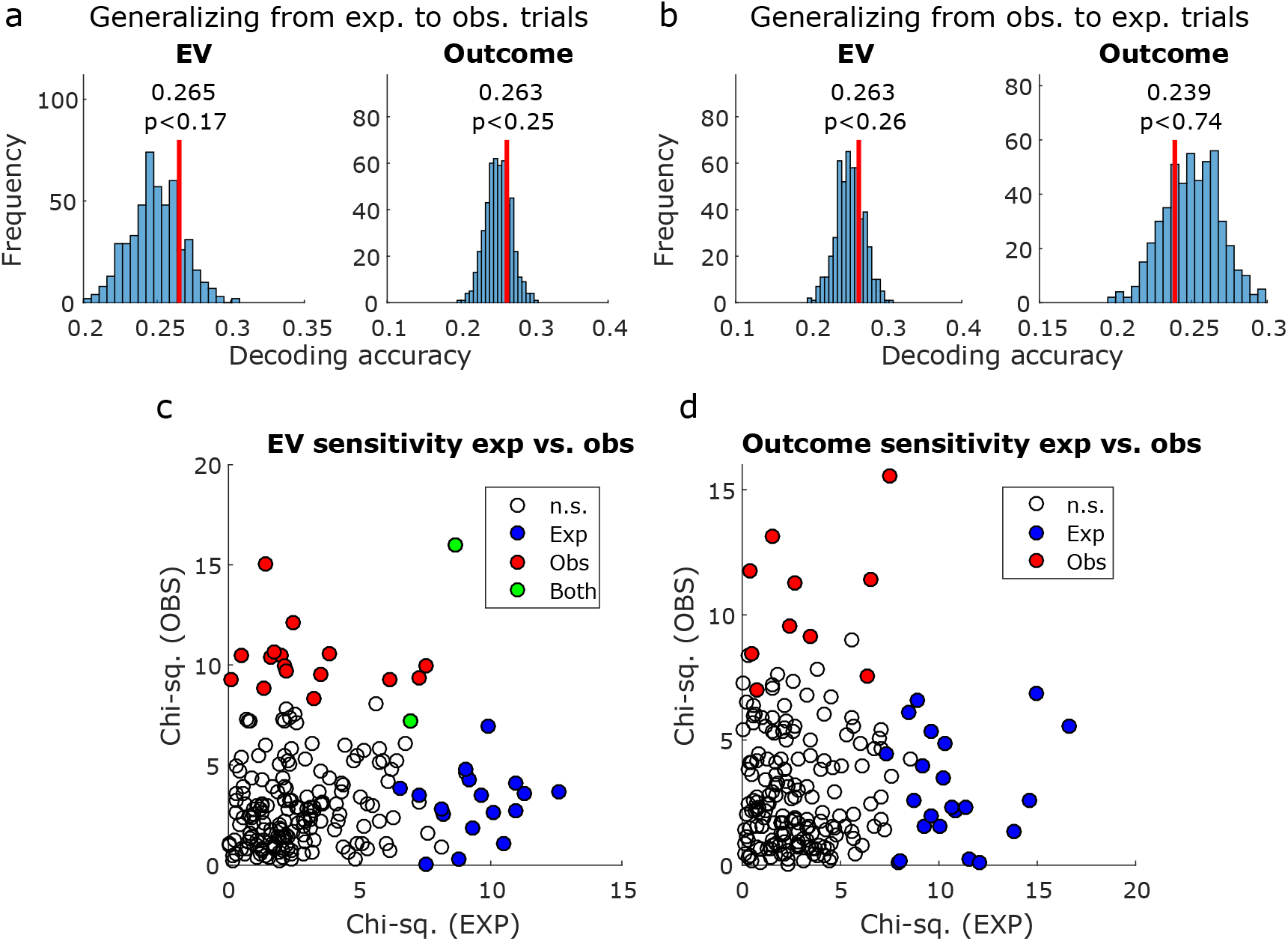
Comparing decoding and encoding across experiential and observational trials. (a) Decoding generalization, training a decoder in experiential trials and testing in observational trials. Decoded variables were EV (left) and outcome (right). Vertical red lines indicate decoding accuracy, and histograms indicate decoding accuracy in each instance of the permutation test with shuffled variable labels. P-values were obtained by computing the proportion of permutation iterations in which the decoding accuracy exceeded the true decoding accuracy. (b) Same, but training in observational trials and testing in experiential trials. (c) Sensitivity to EV in each unit, as obtained in the encoding analysis, plotted for experiential trials (x axis) and observational trials (y axis). Sensitivity was defined as the chi-squared value obtained from the Kruskal-Wallis test used in the encoding analysis. Unfilled data points indicate not sensitive units, blue data points indicate units only sensitive to experiential EV, red data points indicate units only sensitive to observational EV, and green data points indicate units sensitive to both experiential and observational EV. (b) Same, but for outcome sensitivity.

None of the generalization decoding tests yielded better than chance decoding accuracy in the permutation test, regardless of which set of trials (experiential or observational) was used to train or test the decoder.

Additionally, we plotted the sensitivity of each individual amygdala neuron to EVs (pre-outcome, Fig. 7c) and outcomes (post-outcome, Fig. 7d), contrasting experiential and observational trials. The sensitivity of each neuron is defined as the chi-squared value obtained from the previously described encoding Kruskal-Wallis test for differing levels of EV or outcome. The Pearson correlation between EV sensitivities was *ρ* = 0.10 (*p* < 0.14), and only 2 units were found to be sensitive in both trial types. Additionally, the Pearson correlation between outcome sensitivities was *ρ* = 0.02 (*p* < 0.77), and no units were found to be sensitive in both trial types.

These results indicate that despite evidence for successful outcome decoding in each condition separately, and significant unit counts for EV in both conditions, there is no evidence supporting a shared representation between experiential and observational trial conditions in amygdala.

## Discussion

In observational learning, an individual learns about the value of stimuli in the world not through direct experience, but instead through observing the experiences of others. Here we investigated whether the human amygdala contains neuronal representations of key computational variables relevant for learning about the value of stimuli through observation. We found evidence for the encoding of the EV of a stimulus in amygdala neurons, at the time when participants are observing another agent choose that stimulus before this agent received an outcome, even though on those specific trials no tangible reward outcome is obtained by the participant themselves. In addition, we found evidence that the amygdala contains decodable representations of outcomes during observational learning and experiential learning. Together, these results suggest that the human amygdala tracks several key reinforcement learning variables that can be deployed for observational reward-learning.

In addition, human amygdala neurons also strongly discriminated between whether or not a particular trial involved observational or experiential learning at the trial onset. This was by far the most robust signal found in the amygdala neurons, though this could at least in part be an effect of the distinct visual properties of experiential and observational trials (i.e. the presence of the head/face of the observed person, which can modulate amygdala cells (Minxha et al., 2017)), or of the blocked task design. Still, taken together with the RL computations found in the amygdala that were related to observational learning, these findings would appear to support an important contribution of the human amygdala to observational learning.

Consistent with a large literature describing the role of amygdala in anticipating rewards (Belova et al., 2008; Prévost et al., 2013; O’Doherty et al., 2017), we found evidence for experiential EV in both the single unit encoding analysis and the population decoding analysis, as well as for observational EV, in the single unit encoding analysis, further supporting the computational model as a meaningful description of behavior.

An issue that requires further investigation is whether neurons encode experiential and observational expected value signals independently of the identity of the presented stimulus. Previous studies have reported prominent stimulus identity encoding at the single neuron level in amygdala, such as in the recognition of faces and objects (Fried et al., 1997; Gothard et al., 2007), categorization (Kreiman et al., 2000), and visually selective neurons during memory retrieval (Rutishauser et al., 2015b), but also identity-independent stimulus feature encoding, such as in memory-selective neurons during memory retrieval (Rutishauser et al., 2015b) and ambiguity neurons during decision making (Wang et al., 2017).

A related question is whether the neural substrate representing value in amygdala neurons is the same or different for observational and experiential learning. That is, do EVs and outcomes activate amygdala neurons in a similar manner, whether it occurs in an observational learning situation or an experiential learning situation? To test this we trained a classifier to decode these variables in observational learning and tested this classifier on the same neurons during the experiential learning condition and vice-versa. In both cases we could not successfully decode signals when training on one condition and testing on the other. These findings suggest that neuronal coding of observational learning EV and outcomes is distinct and not-overlapping with the neuronal code for experiential learning prediction errors. Additionally, we inspected the sensitivity of individual neurons while encoding EVs and outcomes, and found that across the amygdala neuron population, experiential and observational sensitivities to these variables do not correlate. There is also little overlap between which neurons encode EV and outcomes in each condition. Overall, these findings support a distinction in how these two learning functions are implemented at the neuronal level, at least in the amygdala. These findings also support the argument that the subjects properly understood the task and knew that the observed rewards would not be given to them. If this were not the case, the neural representation for expected values and outcomes likely would not be separate.

It is worth noting that the forward encoding analyses and the backward (decoding) analyses gave slightly divergent results for some of the variables in the observational learning condition. For instance, expected value signals during observational learning were robustly detected in the single-unit encoding analysis but only weakly detectable in the decoding analysis (with the decoding accuracy bordering but not reaching statistical significance). Such divergent results could potentially arise due to differences in the nature of the neural signals being detected by the two methods. The decoding method is liable to be more sensitive to distributed representations across the neuronal population, pooling information from neurons that do not by themselves, individually reach a threshold for significant encoding in the encoding analysis, yet still contain information relevant for representing the variable of interest. On the other hand, the encoding analysis is by definition more sensitive to the role of individual neurons in encoding a particular variable, while being less sensitive to the distributed representation of the variable in question. The minor discrepancies we found between the encoding and decoding analyses in this regard may be informative for better understanding the nature of the neuronal encoding of these different decision variables. Perhaps observational outcome codes are more distributed across the neuronal population in the amygdala, while observational expected value codes are more circumscribed and present in a discrete population of amygdala neurons as opposed to being encoded in a more distributed fashion. Alternatively, these differences might be entirely due to prosaic differences in the statistical methodologies used in the two analysis approaches. Further investigation of observational learning signals in the amygdala will be needed to discriminate these possible explanations.

This notwithstanding, our results highlight the importance of performing population level analyses based on all recorded neurons, as well as the encoding analyses which describe the correlations between the activity of individual units and the variables of interest. Note the population decoding approach can enable a more complete understanding of the information carried by neural population activity even in the presence of mixed selectivity on the level of individual neurons (Rigotti et al., 2013; Saxena and Cunningham, 2019).

In the present study we did not assess whether the observational learning signals we found in the amygdala are specifically recruited when observing another human agent, or rather are recruited when observing causal relationships between stimuli, actions and outcomes irrespective of the nature of the agent performing the actions. An important direction for future studies would be to compare and contrast neuronal effects in the amygdala during observational learning when the agent is human or a computer. There is no strong reason to assume a-priori that the responses detected in the amygdala should be specific only to observed human agents. However, it is possible that the presence of a human might enhance the salience of the observed stimuli compared to the situation where the agent is non-human, which could potentially increase the magnitude of neuronal responses.

To conclude, our findings support an important role for the human amygdala in observational learning, particularly under situations where associations between stimuli and outcomes are learned about through observing the experiences of another agent. The amygdala was found to contain neuronal representations depicting the expected future reward associated with particular stimuli when observing the experiences of another agent interacting with and obtaining rewards from those stimuli. Furthermore, amygdala neurons responded to outcomes whether the subject experienced them or passively observed another agent receiving them. The specific contributions we have uncovered for the amygdala in observational learning adds to a burgeoning literature highlighting a broad role for this structure in social cognition more generally. (Adolphs et al., 1998; Gothard et al., 2007; Adolphs, 2010; Chang et al., 2015; Minxha et al., 2017; Taubert et al., 2018).

## Acknowledgements

This research was supported by NIH (R01DA040011 and R01MH111425 to J.O.D., R01MH110831 to U.R., and P50MH094258 to J.O.D. and U.R.).

## Code availability

The analysis code is available upon request to the authors.

## Author contributions

T.G.A. performed spike data pre-processing and all subsequent data analyses, and wrote the manuscript; J.M. collected data, performed spike data pre-processing and wrote the manuscript; S.D. created the original task design; I.B.R. performed patient surgery; A.N.M. performed patient surgery; U.R. and J.O.D. both conceived and directed the project, and wrote the manuscript.

